# Determining whether a class of random graphs is consistent with an observed contact network

**DOI:** 10.1101/168047

**Authors:** Madhurima Nath, Yihui Ren, Yasamin Khorramzadeh, Stephen Eubank

**Affiliations:** Network Dynamics Simulation and Science Laboratory, Biocomplexity Institute of Virginia Tech, Blacksburg, VA 24061, USA; Department of Physics, Virginia Tech, Blacksburg, VA 24061, USA; Department of Population Health Sciences, Virginia Tech, Blacksburg, VA 24061, USA; Center for Complex Network Research and Department of Physics, Northeastern University, Boston, MA 02115, USA

**Keywords:** Network reliability, Epidemic modeling, Network structure, ERGM, Epidemic potential

## Abstract

We demonstrate a general method to analyze the sensitivity of attack rate in a network model of infectious disease epidemiology to the structure of the network. We use Moore and Shannon’s “network reliability” statistic to measure the epidemic potential of a network. A number of networks are generated using exponential random graph models based on the properties of the contact network structure of one of the Add Health surveys. The expected number of infections on the original Add Health network is significantly different from that on any of the models derived from it. Because individual-level transmissibility and network structure are not separately identifiable parameters given population-level attack rate data it is possible to re-calibrate the transmissibility to fix this difference. However, the temporal behavior of the outbreak remains significantly different. Hence any estimates of the effectiveness of time dependent interventions on one network are unlikely to generalize to the other. Moreover, we show that in one case even a small perturbation to the network spoils the re-calibration. Unfortunately, the set of sufficient statistics for specifying a contact network model is not yet known. Until it is, estimates of the outcome of a dynamical process on a particular network obtained from simulations on a different network are not reliable.

## 1. Introduction

The role of complex networks has become increasingly important in diverse fields of study, ranging from biology to social sciences to engineering. In the field of social sciences, networks often model contacts among a population. The nodes represent the individuals in the population and edges represent the contacts or interactions between them. Applied to infectious disease epidemiology, each edge is associated with a probability of transmitting infection. Simulations draw an instance from the joint probability of infecting any set of people, providing insights into the spread of the disease through the population. Historically, the focus of mathematical epidemiology has been on properties such as the period of incubation, the duration of illness and the mortality rate, and less on the structure of the contact network. Hence simple compartmental approaches that assume a more or less homogeneous mixing have been adequate [1, 2, 3, 4, 5, 6]. However, especially in the context of evaluating targeted control efforts for sexually transmitted diseases, there has been increased emphasis on contact networks.

The transmission of an infectious disease through a contact network can be modeled as a diffusive process on a graph [7, 8, 9]. The size and overwhelming complexity of modern epidemiological problems calls for new approaches and tools like stochastic processes, random walks or Markov Chain Monte Carlo methods. With the aid of computers, agent based models on realistic social networks [9, 10, 11] can bridge from the individual level to population-level. Such models have provided useful insights into the implications of interaction patterns for the spread of disease. These simulations provide a platform to test and understand the spread of diseases and the effects of any intervention measures targeted at specific sub-population [10, 12, 13]. Generalizing results from one region to another requires studying variations in the network and the sensitivity of results to those variations. Because it is difficult to measure large contact networks, these studies rely on drawing sample networks from a network model.

It is known that the structure of the contact network significantly affects the spread of diseases. Even though the behavior of real world systems can sometimes be predicted by random graphs with constraints on structural properties such as degree distribution, discrepancies between theory and simulation, suggest the presence of different social structures which are not captured by these constraints [14]. The effective degree approach [15, 16], edge-based compartmental models [17, 18], modifications to simple compartmental models [19] are a few of the ways researchers have tried to incorporate both the duration of the contacts in the network and the heterogeneities in numbers of partners. In particular, it has been suggested that mathematical models which incorporate such heterogeneities along with the clustering can be used to model the epidemic dynamics on networks [20, 21].

The spread of infectious disease is an example of the classical bond percolation process and it depends on the network structure. For networks whose structure is tree-like, a large class of epidemic models can be solved exactly to provide analytic expressions for the sizes of both epidemic and non-epidemic outbreaks and for the position of the epidemic threshold [22]. For an arbitrary network, the epidemic threshold condition is closely related to the largest eigenvalue of the adjacency matrix, under reasonable approximations [23]. Further, it is shown that the time taken for the epidemic to die out depends on the difference of the two largest eigenvalues of the adjacency matrix [24]. The fluctuations in the connectivity of the network also influence the overall behavior of epidemic spreading by strongly enhancing the incidence of infection [25]. The effects of the *k*-core structure and clustering of the connections on phase transitions have been shown in [26]. Methods like onion decomposition [27] provide insights about the topology around each node allowing the identification of important local structure. The work presented here investigates whether a class of networks with similar local structure exhibits similar dynamics, in particular, the spread of diseases.

A variety of mathematical models are used in the literature to create networks that can imitate the patterns of the links in real networks. Some are random in nature with few parameters fixed whereas others are more structured and take into account more network properties. Methods such as preferential attachment [28] can generate networks with a certain degree distribution while small world models reproduce the clustering in observed networks [29]. Exponential random graphs [30, 31] create a network model with maximum entropy consistent with matching user-specified properties. Networks drawn from this model have values of these statistics that are closely fitted to those of an observed network, but otherwise random [31]. Recent studies have used the exponential random graph models (ERGMs) to model friendship networks [32]. For example, the spread of sexually transmitted disease has been studied extensively [33, 34, 35, 36, 37, 38] using data available from Wave I of the National Longitudinal Study of Adolescent to Adult Health (Add Health). Add Health is a longitudinal study of a nationally representative sample of more than 90,000 adolescents in grades 7 through 12 in the United States, obtained from the data collected between 1994 and 1995 through a stratified sample of 80 schools. This survey data combines the different demographic factors with the social interactions of these school students. They have been analyzed by Resnick et. al. [39] and Udry and Bearman [40], for example, to identify the characteristics associated with health and risky behaviors among the adolescents. ERGMs generate networks efficiently when the constraints are placed only on local statistics like degree distribution (number of people connected to a certain individual), clustering or number of triangles. But the spread of a disease depends on global properties of the network, which are more difficult to match with an ERGM.

It is imperative to understand how the choice of properties to constrain affects the simulated spread of disease. This paper applies the concept of network reliability, introduced by Moore and Shannon [41] to characterize the effects of the network model. The network reliability *R*(*x*; *α*) takes into account the structural properties (i.e., topology) as well as the dynamics of contagion on the network. It gives the probability of observing an infection attack rate of at least *α* for an SIR (susceptible-infected-recovered) process with transmission probability *x* on a particular network. This is a measure of the “epidemic potential” introduced by Hamilton [42]. Since ERGMs are thought to capture the structural features of social networks [30, 32, 43, 44], the network reliability of an ensemble of ERGMs intended to represent a specific network in the Add Health survey is evaluated. It is observed that even though these models are a good representation of local structure in the network, they lead to significantly different dynamics for the propagation of an epidemic. However, because the transmission probability and the network structure are not separately identifiable, there is a simple transformation of the transmission probability that can erase these differences. This suggests supplementing the ERGM model with a description of this transformation to arrive at an ensemble of social network plus the probability of transmission for simulating the spread of disease. The effect of an intervention measure like vaccination (represented as node removal) is briefly discussed for the networks. The transformation method works well for estimating the overall attack rate even for the networks where the nodes are removed. Unfortunately, a model calibrated to reproduce the overall attack rate does not necessarily reproduce the full time dependence of the epidemic curve, and thus is not well suited for estimating the effects of time-dependent control measures.

## 2. Methods

One of the Add Health friendship networks is chosen as the “observed” network for this study. The results on this population-based survey are compared to those obtained from networks created using the ERGM [45] model based on the observed network. One of the ERGM models yields the well-known Faux Magnolia dataset [45, 46, 47]. The Faux Magnolia network matches the Add Health data in degree distribution, clustering, number of triangles and other centrality measures, indicating that the ERGM is a good candidate model for a social network. The population-based data used for this study is obtained from Wave I of the Add Health study (http://www.cpc.unc.edu/projects/addhealth). One of the friendship networks, school 86 (based on the schools 086 and 186, a junior and a senior high school) is used as the original dataset. A network containing the mutual friends is considered for this study. The ERGMs are used to model the underlying structure of the friendship network. There are different ERGMs available in the statnet package [45] depending on the property to be constrained. Four distinct ERGMs are used to generate four sets of networks, each containing an ensemble of 100 networks. Each of these four sets match the features of the original network, e.g., the total number of edges, node attributes and different values of the GWESP (geometrically weighted edgewise shared partner) statistic [45], a parameter that combines the clustering and the number of triangles in the networks [30, 44]. The details of these models are mentioned in the Section 2.1.

The Faux Magnolia network extracted from the statnet package [45, 46, 47] is an ERGM fit based on this Add Health data. Model 1 constrains the total number of edges of the original dataset; model 2 constrains the node attributes; model 3 constrains the total number of edges and the number of triangles; model 4 constrains the total number of edges, the node attributes and an additional statistic called GWESP [30, 44, 45], which is related to the number of triangles and clustering in a network. The networks in the last two sets are generated using this model, with two different values of the GWESP statistic, 0.25 and 0.5. These models are named to be consistent with the *statnet* naming convention.

### 2.1. Network Generation

The exponential random graph model (ERGM) is used to generate networks with characteristics similar to a friendship network from the Add Health survey. The original network is one that has been built from the survey data of school 86 containing only the mutual interactions. The different ERGM models used for this study are available in the *statnet* package[45, 48]. Four sets of ERGM networks, each set containing an ensemble of 100 networks are generated using the three models. Model 1, referred to here as “edges”, takes the total number of edges of the original data as the constraint. Model 2, referred to here as “node attributes”, uses the node attributes like race, sex and grade along with the total number of edges of the original network to generate the ERGM fits. Model 3, which constrains the number of edges and the number of triangles, failed to converge in the trials [45, 48], so it is not reported here. The last model is further constrained and is used for the remaining two sets. The networks generated using this model use two different values of the GWESP (geometrically weighted edgewise shared partner) statistic [30, 44, 45, 48], 0.25 for model 4.1, referred to here as “GWESP = 0.25”, and 0.5 for model 4.2, referred to here as “GWESP = 0.5”. GWESP is a parameter that affects the clustering and the number of triangles in the networks. The following steps are used to build these networks. Here, *sch* is a R network object that contains the mutual edges from the junior and senior high schools, school 086 and school 186.

~~~
model1 = ergm(sch∼edges)

model2 = ergm(sch~edges + nodematch(“grade“)
                 + nodematch(“race“) + nodematch(“sex“))
model3 = ergm(sch~edges + triangles, verbose=TRUE, maxit = 25,
                  control = control.ergm(steplength = 0.2))
model4_1 = ergm(sch~edges + absdiff(“grade“)
                  + nodematch(“grade“) + nodematch(“race“)
                  + nodematch(“sex“) + gwesp(0.25, fixed=TRUE),
                  burnin = 1e+4, interval = 1000,
                        MCMCsamplesize = 2500, maxit = 25,
                 control = control.ergm(steplength = 0.25))
model4.1 = simulate(model4 1, burnin = 1e+8, constraints=∼edges)

model4 2 = ergm(sch~edges + nodematch(“grade“)
                    + nodematch(“race“) + nodematch(“sex“)
                    + gwesp(0.5, fixed = TRUE), MCMCsamplesize = 1e+5,
                    maxit = 15, verbose=TRUE,
                         control = control.ergm(steplength = 0.25))
model4.2 = simulate(model4 2, burnin = 1e+6, verbose=TRUE)
~~~

The Faux Magnolia network is extracted from the *statnet* package [45]. “The Faux Magnolia data set represents a simulation of an in-school friendship network. It is based upon the schools 086 and 186 from the Add Health Wave I dataset.” [48] Table 1 summarizes model constraints.

**Table 1:** Summary of constraints used in [45, 48] for constructing the ERGM models.

| *statnet* Model | Constraints Used | Labels Used |
| --- | --- | --- |
| model 1 | Total number of edges | edges |
| model 2 | Edges + Node attributes | node attributes |
| model 3 | Edges + Triangles |  |
| model 4.1 | Edges + Node attributes + GWESP = 0.25 | GWESP = 0.25 |
| model 4.2 | Edges + Node attributes + GWESP = 0.5 | GWESP = 0.5 |

### 2.2. Statistics on Networks

Typical measures of network structure like degree distribution, number of triangles, clustering coefficients and centrality measures closeness and betweenness centrality are calculated for all the networks generated by the different models as well as the Faux Magnolia and the school 86 networks (Supplementary Notes). Comparison of these measurements demonstrates that the final model, “GWESP = 0.5”, and the Faux Magnolia network are best calibrated while all the models meet the constraints to the other statistics as well.

### 2.3. Epidemic Threshold for the Networks

The epidemic threshold condition for networks which have tree-like structure locally given by Newman’s formula [22] can be written in terms of the mean *k* and the variance *Var*[*k*] of the degree distribution [15, 16]. This quantity, called *x_c_* in this paper, is given by the Equation 1.

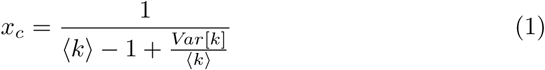

For an arbitrary graph, *x_c_* is inversely related to the largest eigenvalue of the adjacency matrix [23]. The values of the largest eigenvalues for these networks are similar, *λ*_86_ = 5.05 and *λ_FM_* = 4.98. The Figures 1a and 1b show that *x_c_* for the school 86 network and the Faux Magnolia are similar compared to the other network models. (Plots for the largest eigenvalues and the difference of the two leading eigenvalues are in the Supplementary Notes.) The boxplots indicate that Faux Magnolia matches the *x_c_* values much better than the others. It is to be noted that the networks obtained using model “edges” and model “node attributes” have a higher threshold value in contrast to the models “GWESP = 0.25” and “GWESP = 0.5”.

**Figure 1:**
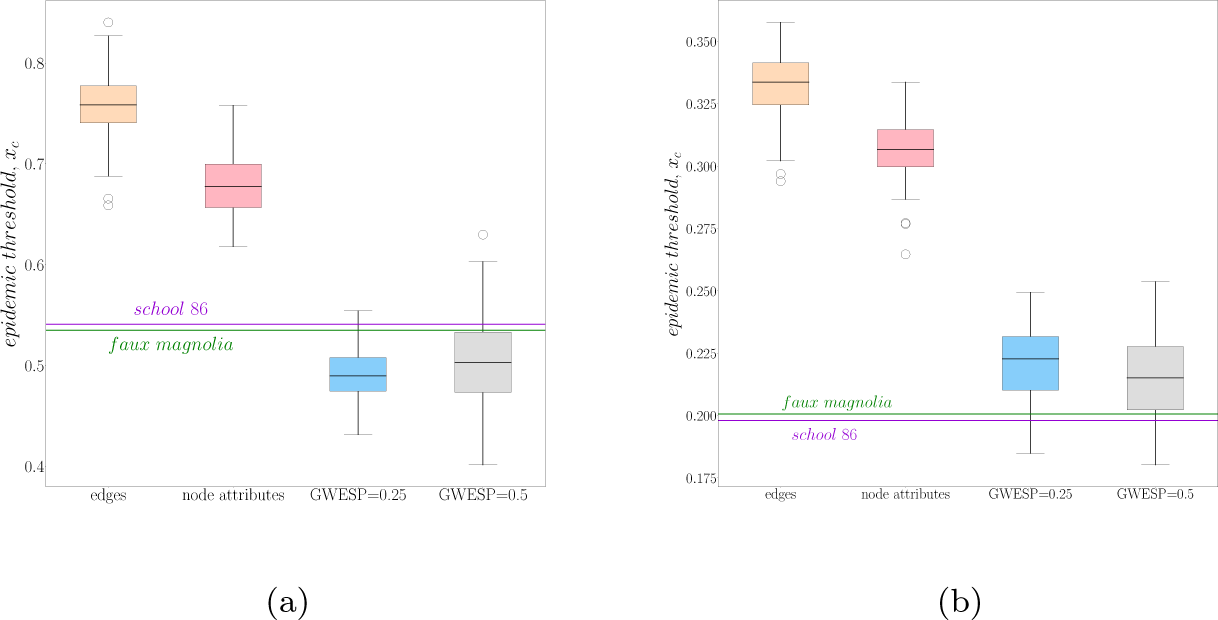
(Color online) Boxplots showing the values of the epidemic threshold, *x_c_* for the networks. The threshold is calculated using Newman’s formula in (a) and using the largest eigenvalue of the adjacency matrix in (b). The labels on the horizontal axis correspond to the models used.

The mean epicurves obtained after 10^4^ SIR simulations for all the 100 networks for each model with those for school 86 and Faux Magnolia are shown in Figure 2. The plots in Figure 3a use one randomly chosen network from all the models for the simulation. The error bars represent the probable errors for the estimated mean value. It can be concluded that the epicurve for Faux Magnolia is the best match for school 86 network. However, despite the similarity in the shape of the epicurves of the two networks in Figure 3b, there is a systematic difference between them. The curve corresponding to Faux Magnolia overestimates the length of an outbreak and the height of the peaks for a particular value of the transmission probability. Further, the average attack rate defined as the average of the total number of the people infected when a randomly chosen individual is infected with the given values of *x* − is ~ 0.0651 for Faux Magnolia and ~ 0.0402 for school 86 when *x* = 0.85. The attack rates for networks obtained from the models “edges”, “node attributes”, “GWESP = 0.25” and “GWESP = 0.5” are ~ 0.1308, 0.2212, 0.2634 and 0.2425 respectively.

**Figure 2:**
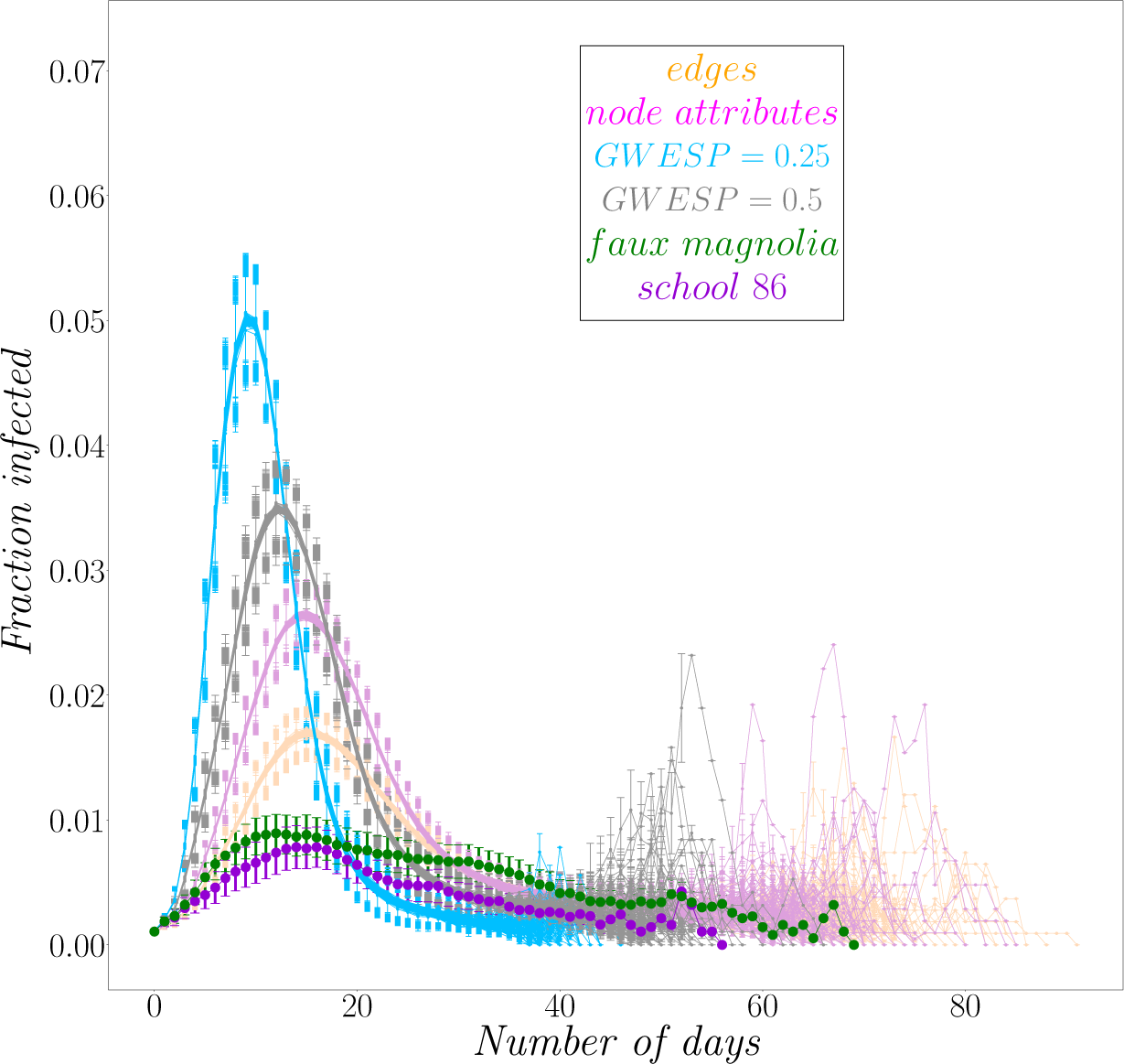
(Color online) Summary of epicurves for all the networks for the probability of infection = 0.85.

**Figure 3:**
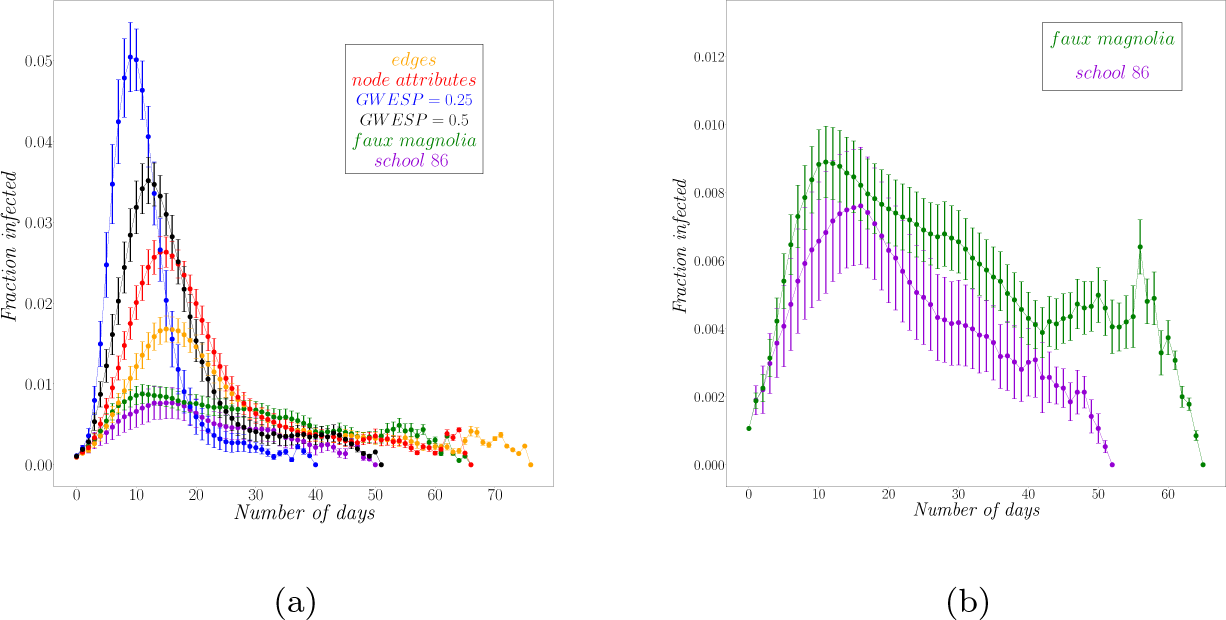
(Color online) Mean epicurves for the networks for a probability of transmission, *x* = 0.85. The error bars are the probable errors for the estimated mean value. (a) This figure shows the mean epicurves for one of the networks from each of the models along with the Faux Magnolia and school 86 networks. (b) This detail from (a) shows the epicurves for the Faux Magnolia and school 86 networks.

### 2.4. R(*x;α* Epidemic Potential (Network Reliability)

*R*(*x;α*), the probability of observing an attack rate of at least *α* for an SIR process with transmission probability *x* on a particular network is estimated [49]. For example, *R*(*x*; *α* = 0.3) gives the probability that the overall attack rate for an SIR process is at least 30% when the probability of transmission of infection is *x*. This measure depends on both the network structure and *x*. Thus, it specifically reflects the behavior of SIR dynamics on a particular network. It is evaluated for all the networks for three different values of *α*, 0.02, 0.05 and 0.08. This is identical to calculating the probability that at least 2%, 5% or 8% of the population is infected. Simulations are used to verify that the method based on the reliability statistic estimates the correct value of the probability of a disease outbreak. Few values of the transmission probability obtained by the transformation are used as the infection rate to calculate the overall average attack rate in these two networks. The table in the Supplementary Notes shows the results of SIR simulations on the Faux Magnolia network and the school 86 network for different values of *x*.

It is observed that different networks have the same value of *R*(*x*; *α*) for different values of *x* (Figure 5). Taking advantage of this confounding, one model can be calibrated to another, so that the epidemic potential [42], *R*(*x*; *α*) is the same for both. A low order polynomial transformation provides a good fit for the calibration curve. For this analysis, the Faux Magnolia network, school 86 network and one of the networks from model “node attributes”, named net1 in the paper are used. Further, to validate the results obtained, SIR simulations are run on these networks for different values of *x* and the values of the average attack rates are compared.

Figure 4a shows the values of *R*(*x*; *α*) for the Faux Magnolia and the school 86 networks for the three values of *α* mentioned above. The scatter plot in Figure 4b shows the variation of the overall attack rates for both the networks as a function of the attack rate for school 86 for all values of probability of transmission *x*. Figure 6 shows the values of *R*(*x*; *α*) for the different networks over a range of all the possible values of *x* for *α* = 0.05. Each shaded region with a curve showing the median value represents each ensemble, and the shaded area lies between the 5*^th^* and the 95*^th^* quantile curves. (Results for other values of *α* are in the Supplementary Notes.)

**Figure 4:**
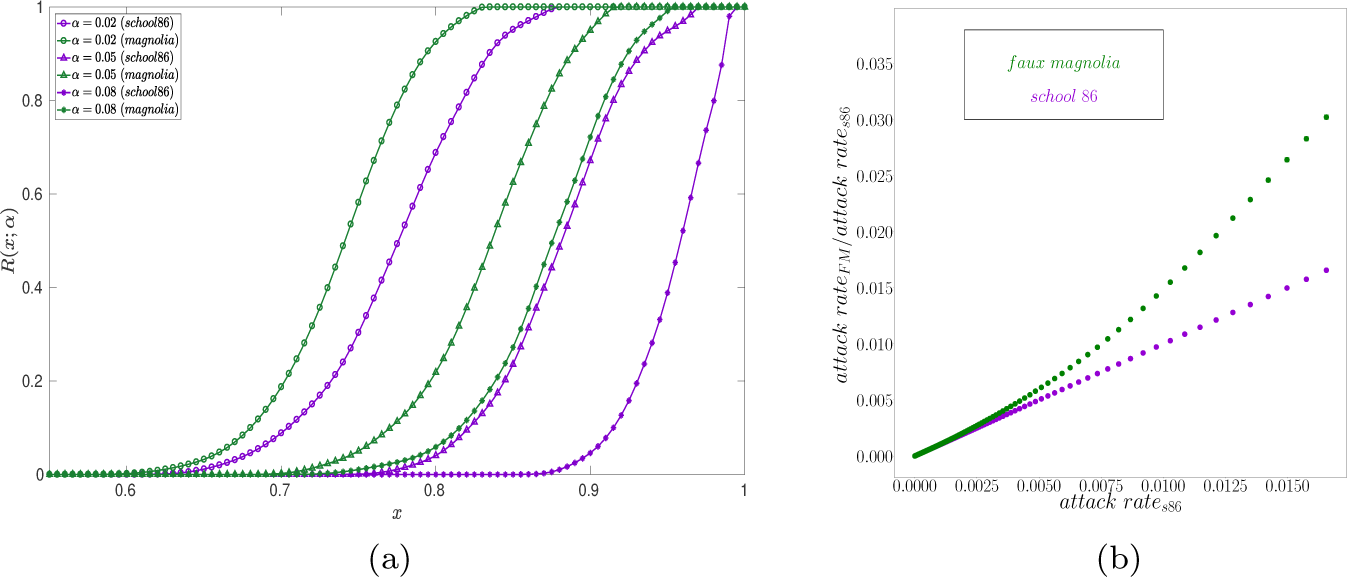
(Color online) (a) *R*(*x*; *α*) for school 86 and Faux Magnolia networks for three values of the attack rates. (b) The overall attack rate for school 86 and Faux Magnolia as a function of the attack rate for school 86. The error bars (which are of the size of the points) represent the mean probable error in estimating the attack rate after 1000 simulations.

From Figure 6, it is evident that different values of the transmission probability, *x* correspond to the same value of *R*(*x*; *α*). Figure 5 shows a schematic of how *R*(*x*; *α*) for two different networks, G1 and G2 are the same for the *x* and *x_eff_* . This property can be used to generate a re-calibration curve. Empirically, it turns out that the two values of *x* are related to each other by a quadratic (details in the Supplementary Notes). The transformed *x* values of the Faux Magnolia network relative to those for the school 86 network are estimated using a quadratic fit, i.e.,

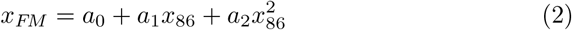

**Figure 5:**
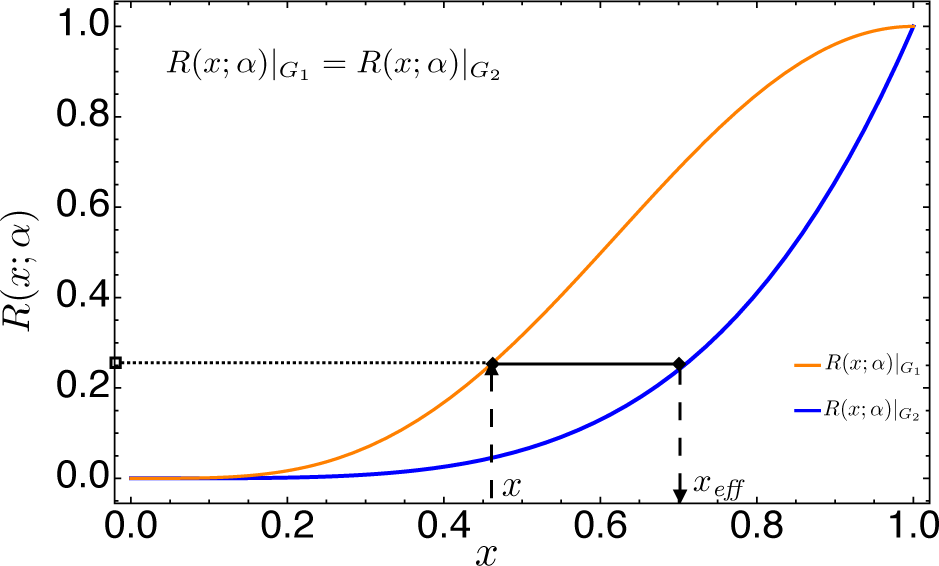
(Color online) *R*(*x*; *α*) has the same value for the networks G1 and G2 at *x* and *x_eff_*.

**Figure 6:**
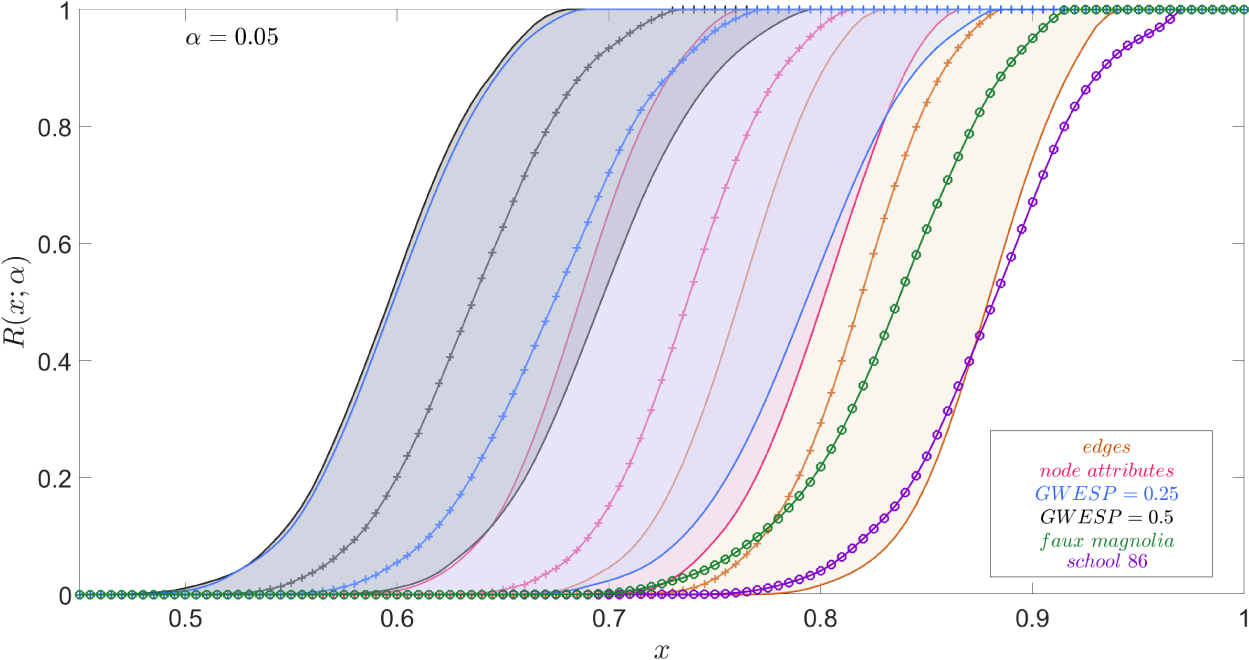
(Color online) *R*(*x*; *α*) values for all networks for a range of values of *x*. Each shaded region represents one of the four models with the crossed lines representing the medians for each set and the solid lines denoting the 5*^th^* and the 95*^th^* quantile curves. The lines with the circles display the values for Faux Magnolia and school 86 networks.

The *x* values re-calibrated according to Equation 2 reproduce the epidemic potential for the networks. The plot in Figure 7 shows these estimated values of *R*(*x*; *α*) obtained using the quadratic polynomial fit from the transformed values of *x* for the case when *α* = 0.05. (The plots for *α* = 0.02 and 0.08 are in the Supplementary Notes.) It is to be noted that the estimated values calculated using this technique are as good as those calculated numerically for these two networks.

**Figure 7:**
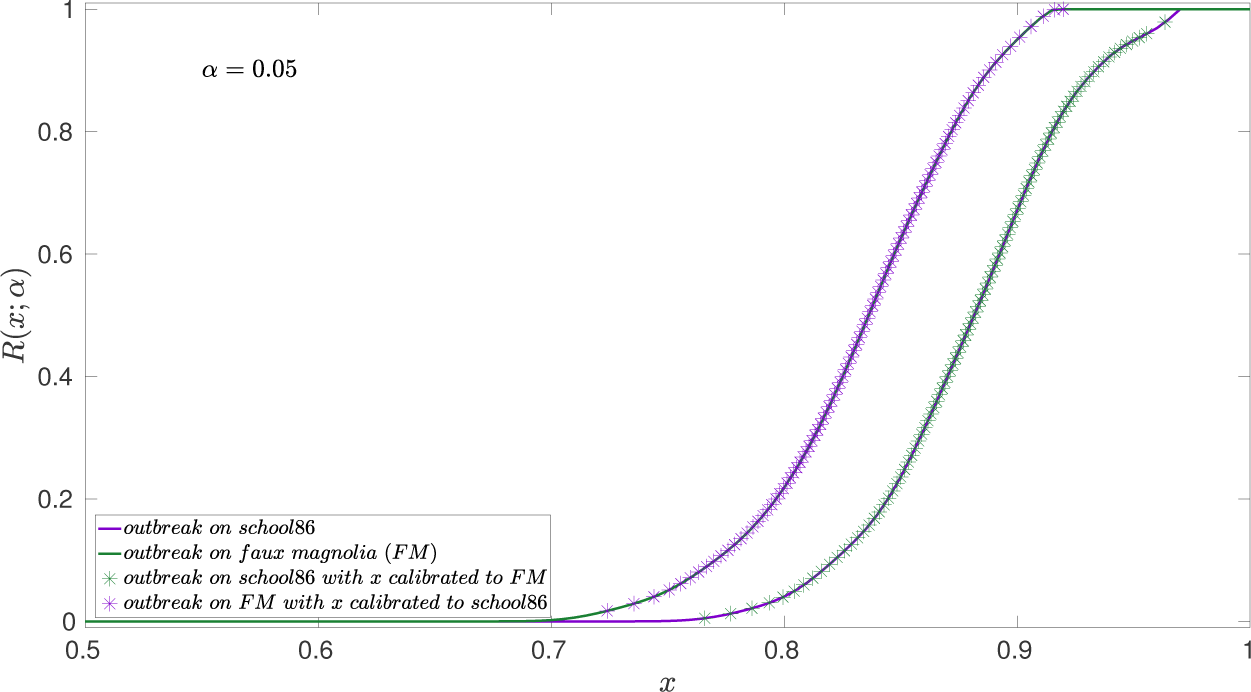
(Color online) Plot showing that the estimated values of *R*(*x*; *α*) for the Faux Magnolia network calculated using the quadratic polynomial fit from the values obtained from the school 86 network agree with those obtained from the numerical analysis on the Faux Magnolia network.

Figure 8 shows the time evolution of the overall fraction of infected people for the two networks Faux Magnolia and school 86 - for two values of the probability of transmission, *x* = 0.85 and 0.92. These are the mean epicurves obtained from 10^4^ SIR simulations on these networks. The error bars represent the probable error for the estimate of the mean value from the simulations. The epicurves for each run of the simulations and their mean curve are presented in the Supplementary Notes. These two networks are further compared with net1 (the network obtained from the model “node attributes”). The mean epicurves for all three networks are plotted in Figures 8a and 8c for the above values of *x*.

**Figure 8:**
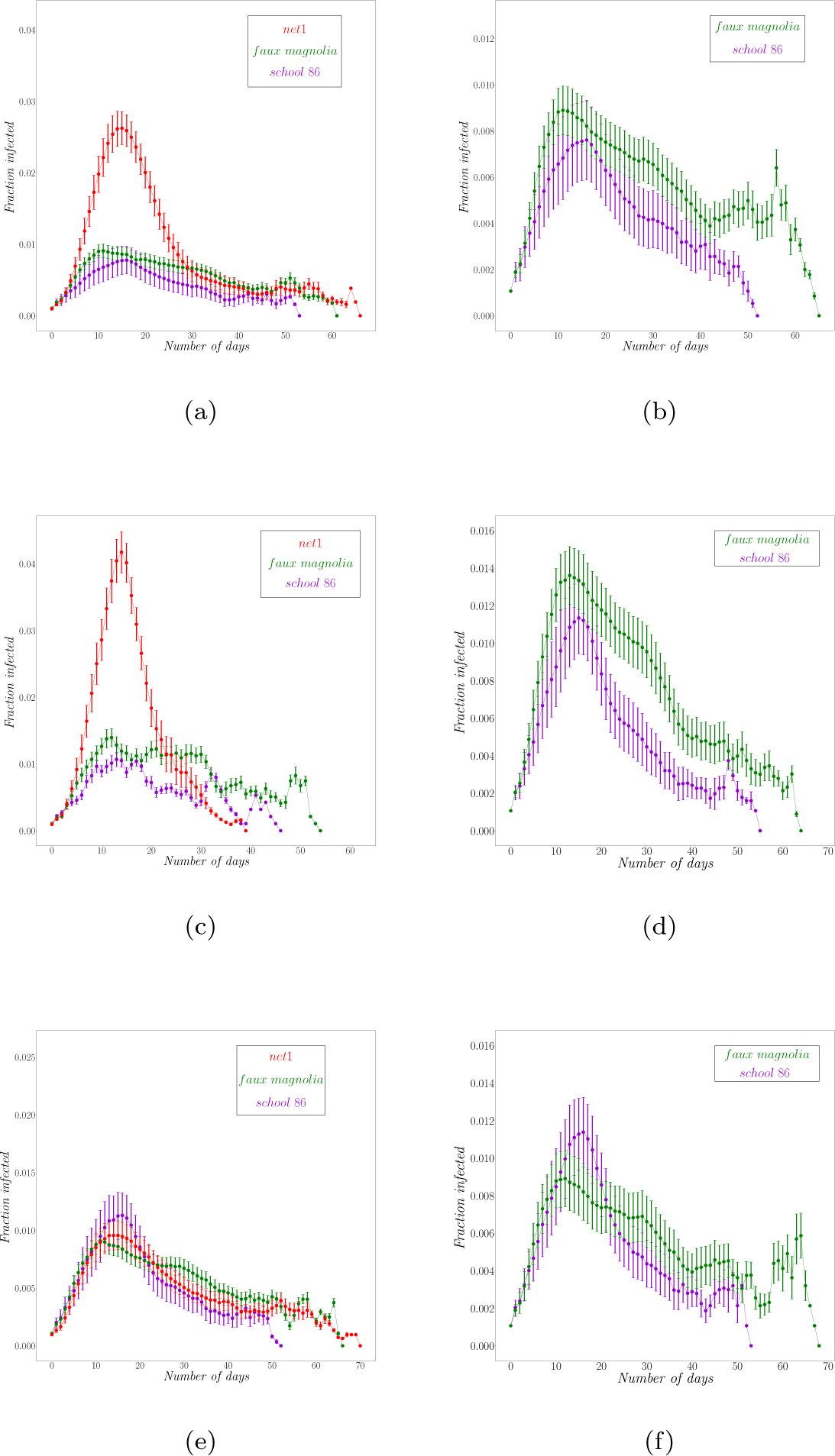
(Color online) The mean epicurves for school 86 and Faux Magnolia networks for two values of the probability of transmission, *x*. (a) and (b) when *x* = 0.85 and (c) and (d) when *x* = 0.92. (a) and (c) show the epicurves for these networks along with net1. (e) and (f) The mean epicurves for the networks for three values of transmission, *x* = 0.73 for net1, *x* = 0.85 for Faux Magnolia and *x* = 0.92 for school 86, to obtain the same average attack rate (~ 0.06). The error bars are the probable errors for the estimated mean value.

The re-calibration is done to obtain an overall attack rate of ~ 0.06 for all three networks. Plots 8e and 8f show the mean epicurves for different values of *x* chosen so that the average attack rate is similar. Figures in the right panel, i.e, 8b, 8d and 8f allow a detailed comparison between school 86 and Faux Magnolia. The overall attack rate of ~ 0.06 is obtained in the school 86 network for *x* = 0.92 whereas in the Faux Magnolia network it is obtained when *x* = 0.85 (also shown in the table in the Supplementary Notes). For the random network (net1), a lower value, *x* = 0.73 gives the same attack rate.

### 2.5. Effects of Intervention Measures

To investigate the response to intervention measures some nodes are removed at random from each of the original networks to capture the effects of vaccination. Two new networks are generated from each of school 86, Faux Magnolia and net1 networks by removing 10 and 100 nodes at random. Figures 9 and 10 compare the mean epicurves obtained after 10^4^ SIR simulations on the new networks for two values of transmissibility, *x* = 0.85 and 0.92. The error bars represent the probable error in estimating the mean curves. In both the figures, the plots in the right panel allow a detailed comparison between school 86 and Faux Magnolia.

**Figure 9:**
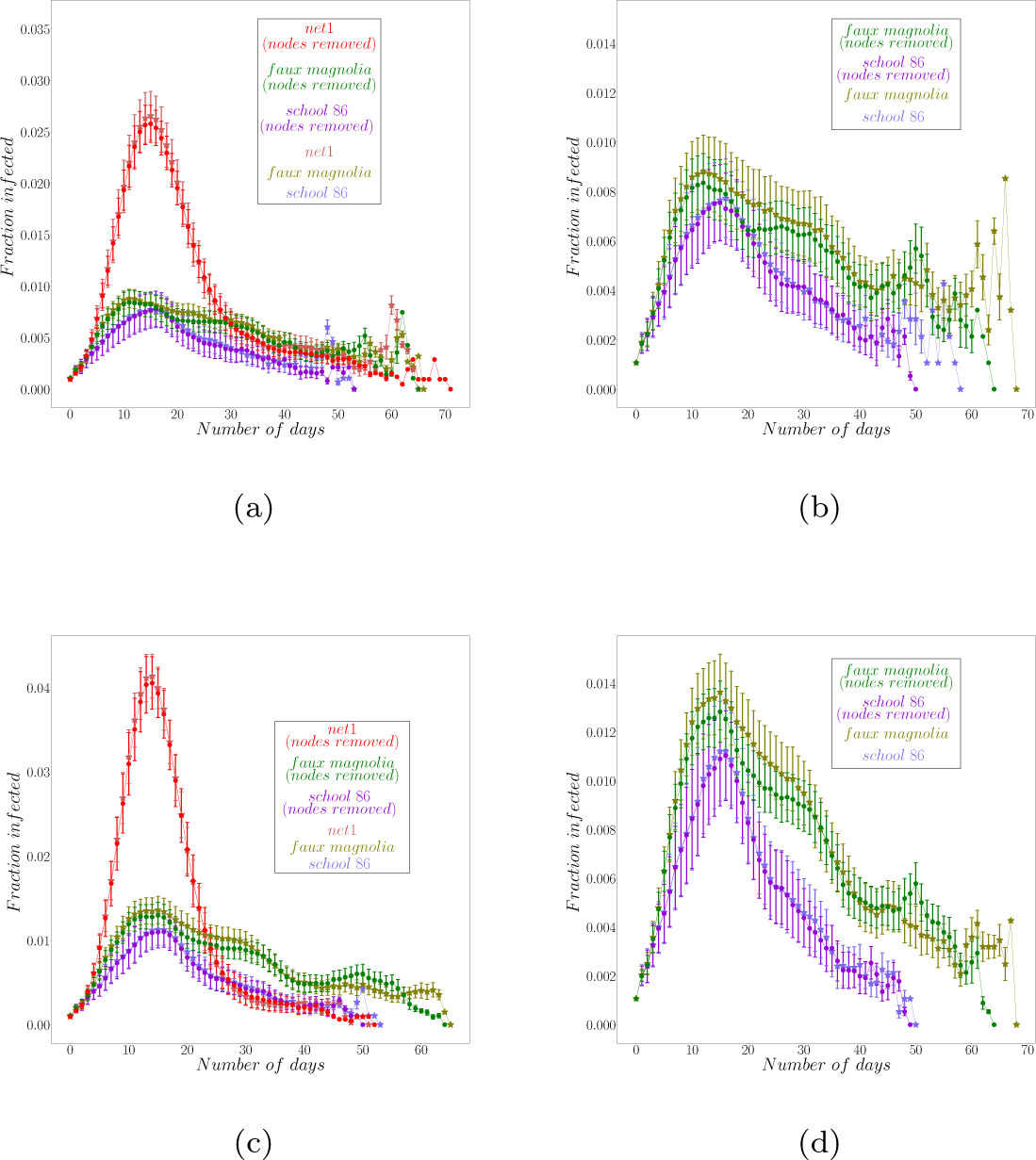
(Color online) The mean epicurves for school 86, Faux Magnolia and net1 networks for two values of the probability of transmission, *x*. (a) and (b) when *x* = 0.85 and (c) and (d) when *x* = 0.92. (b) and (d) show the detailed epicurves for school 86 and Faux Magnolia. The error bars are the probable errors for the estimated mean value. The starred curves represent the original networks; the dotted curves, networks with 10 nodes removed.

**Figure 10:**
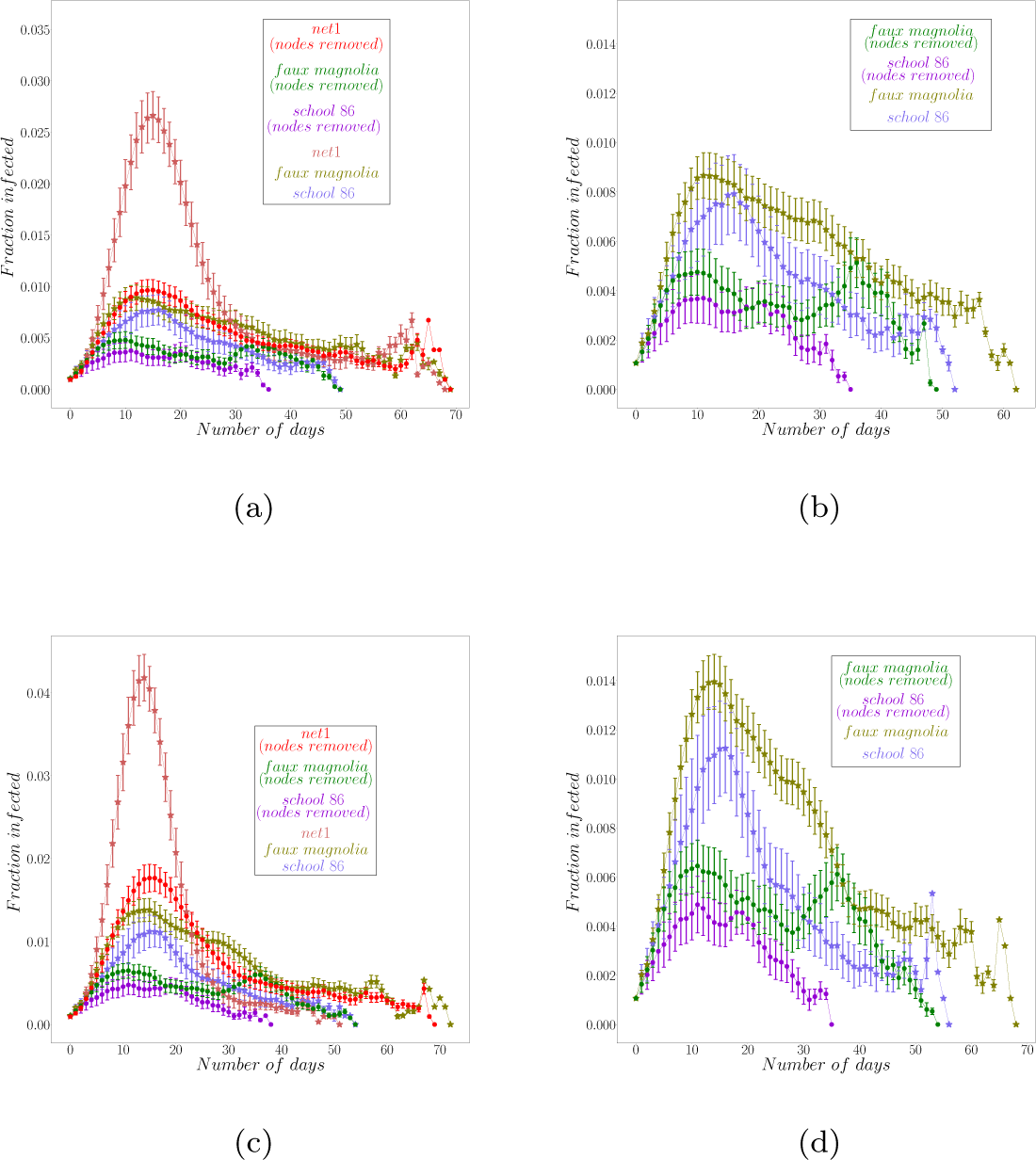
(Color online) The mean epicurves for school 86, Faux Magnolia and net1 networks for two values of the probability of transmission, *x*. (a) and (b) when *x* = 0.85 and (c) and (d) when *x* = 0.92. (b) and (d) show the detailed epicurves for school 86 and Faux Magnolia networks. The error bars are the probable errors for the estimated mean value. The starred curves represent the original networks; the dotted curves, networks with 100 nodes removed.

The *R*(*x*; *α*) curves for the networks with 10 nodes removed are shown in Figure 11. Figure 12 shows the transformation of the probability of transmission given for the original networks as well as the one obtained by node removal for *α* = 0.08. The *x* values for the networks before and after the removal of the nodes are plotted by the red and the green curves respectively. The plots for *α* = 0.02 and 0.05 are in the Supplementary Notes.

**Figure 11:**
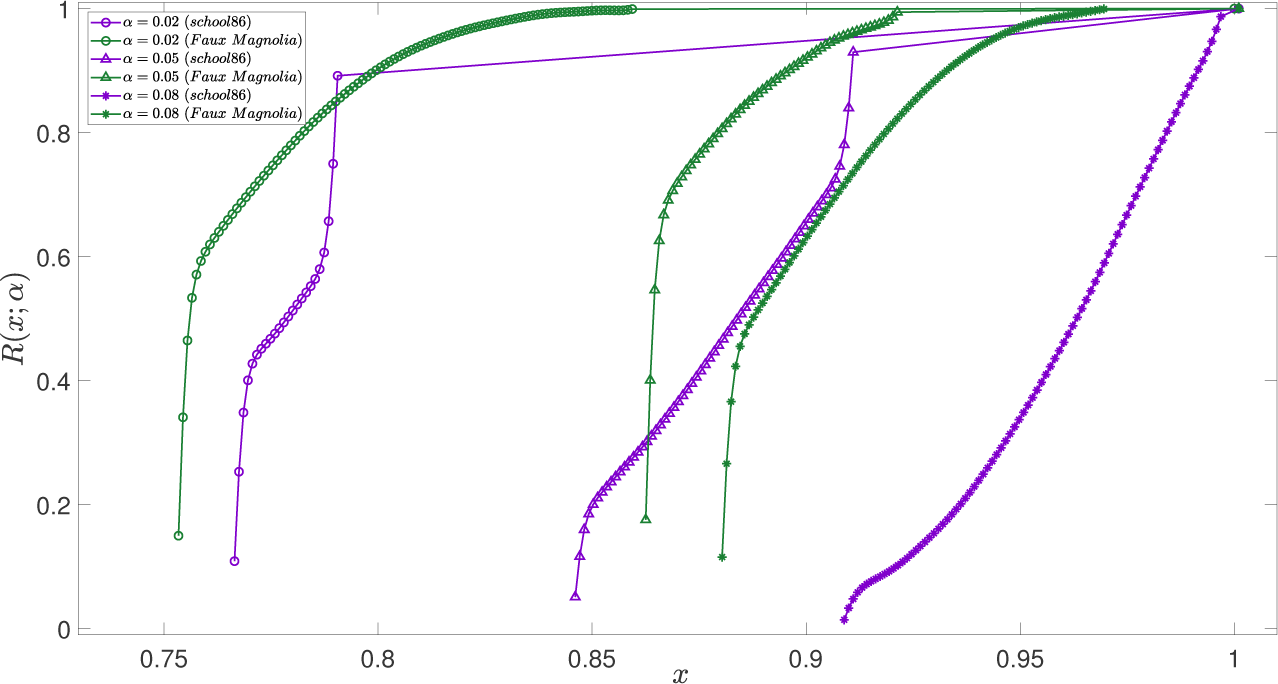
(Color online) *R*(*x*; *α*) for school 86 and Faux Magnolia networks when 10 nodes have been removed from them at random. The x-axis is expanded to provide a better view of the ‘interesting’ region.

**Figure 12:**
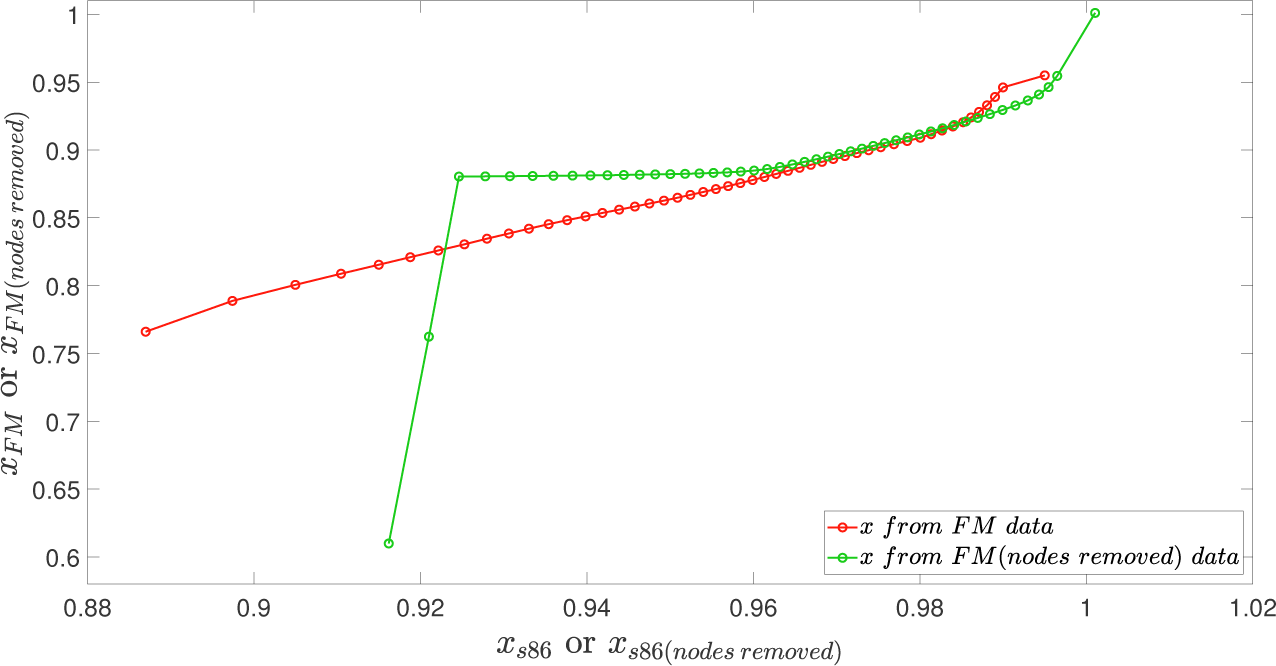
(Color online) Plots showing the *x*-values for Faux Magnolia obtained from the school 86 for *α* = 0.08. The red line represent the transformation for the original networks, the green line with 10 nodes removed. Figures 13 exhibit that the transformation when nodes are removed from the networks. The epicurves for the three networks when 10 nodes are removed 310 are plotted in Figure 13a and 13b and those when 100 nodes are removed are plotted in 13c and 13d. Table 2 summarizes these values for each network. To get a detailed contrast between school 86 and Faux Magnolia networks, their results are presented in the plots in the right panel.

**Figure 13:**
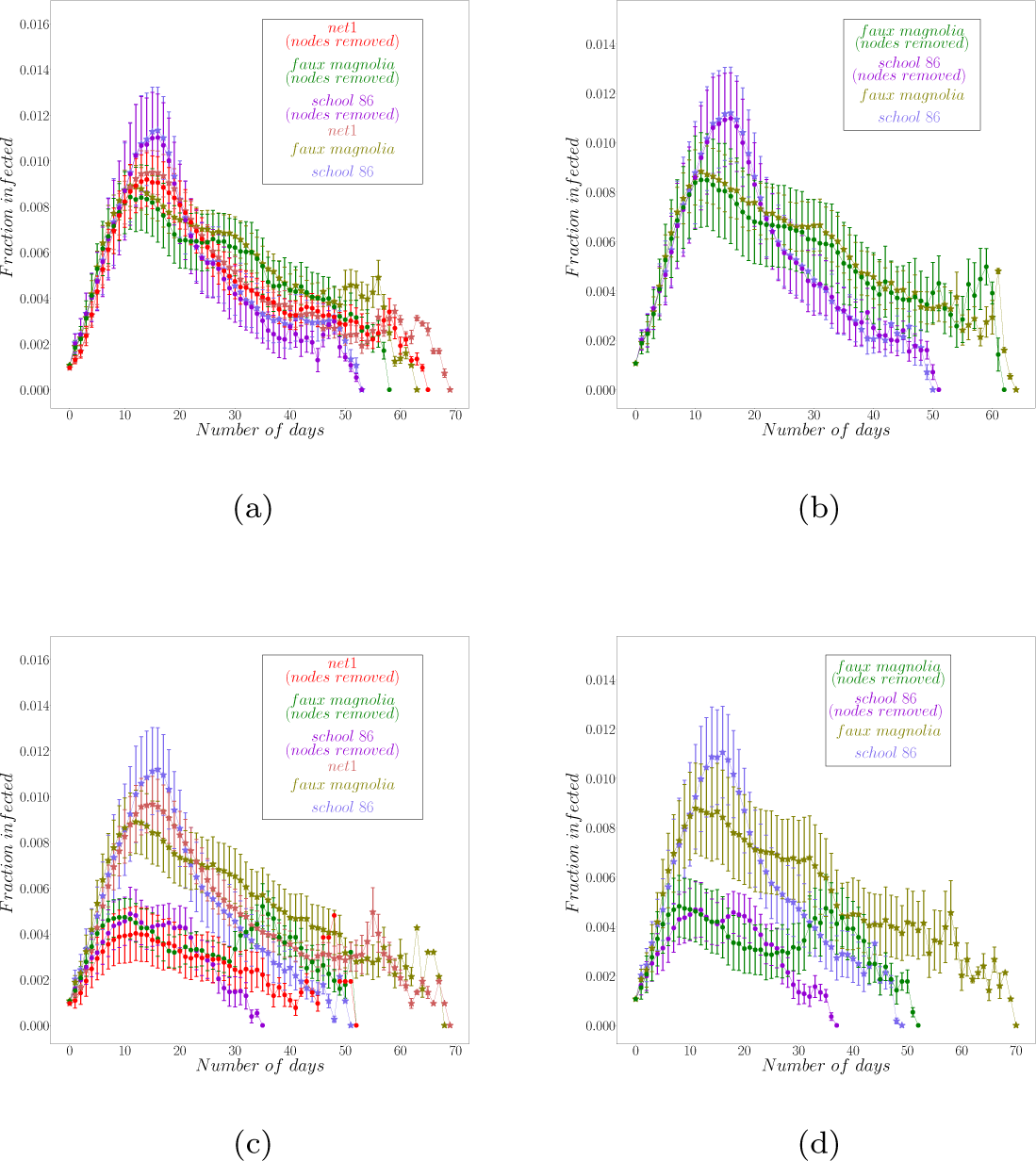
(Color online) The mean epicurves for school 86, Faux Magnolia and net1 for different values of *x* when nodes are removed from the orginial networks. (a) and (b) 10 nodes removed, (c) and (d) 100 nodes removed. The error bars are the probable errors for the estimated mean value. The starred curves represent the original networks; dotted curves, networks with nodes removed.

**Table 2:**
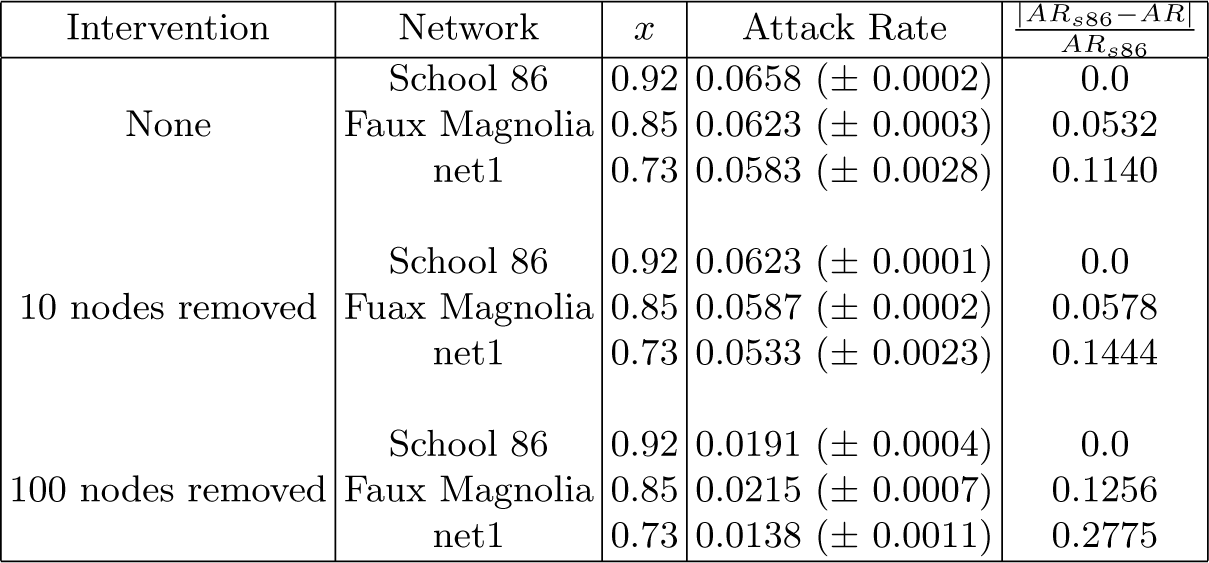
Attack rates obtained for different values of probability of transmission, *x*, for three networks.

| Intervention | Network | $x$ | Attack Rate | $\frac{\vert AR_{s86} - AR\vert }{AR_{s86}}$ |
| --- | --- | --- | --- | --- |
| None | School 86 | 0.92 | 0.0658 (± 0.0002) | 0.0 |
| | Faux Magnolia | 0.85 | 0.0623 (± 0.0003) | 0.0532 |
| | net1 | 0.73 | 0.0583 (± 0.0028) | 0.1140 |
| 10 nodes removed | School 86 | 0.92 | 0.0623 (± 0.0001) | 0.0 |
| | Faux Magnolia | 0.85 | 0.0587 (± 0.0002) | 0.0578 |
| | net1 | 0.73 | 0.0533 (± 0.0023) | 0.1444 |
| 100 nodes removed | School 86 | 0.92 | 0.0191 (± 0.0004) | 0.0 |
| | Faux Magnolia | 0.85 | 0.0215 (± 0.0007) | 0.1256 |
| | net1 | 0.73 | 0.0138 (± 0.0011) | 0.2775 |

## 3. Discussion

For epidemiological purposes, it is essential to explore the effects of the network structure on the dynamical process of an infectious disease epidemic. The boxplots in Figure 1 indicate that although the ERGM succeeds in generating random graphs with similar degree distribution and clustering, uncontrolled variation in other structural properties creates significant change in the value of the epidemic threshold, *x_c_*. The difference in the threshold values in the plots 1a and 1b could be because of the approximation of the tree-like structure in the Newman’s formula [22]. Even though this is an important measure to study the spread of an infectious disease, it does not capture all of the dynamics. The epicurves (Figure 3) provide a better understanding of the whole process. The similarity of epicurves obtained from the SIR simulations implies that the additional constraints used to generate the Faux Magnolia help predict the time evolution of a disease on the school 86 network.

Instead of determining how the system behaves for a particular value of the probability of transmission, a measure that estimates the size of an epidemic outbreak for all its possible values, *R*(*x*; *α*) is suggested [49]. Comparing *R*(*x*; *α*) among different instances of ERGMs shows that, although the Faux Magnolia network model matches the desired statistics of the school 86 network better than the others, the behavior of the disease outbreak is significantly different. Figure 4a shows that the original network is significantly more resistant to disease spread than Faux Magnolia, even though they have similar local statistics. Figure 6 suggests that the school network is more resistant to an epidemic than ERGMs derived from it, i.e., a higher transmission probability is required for widespread disease in the school 86 compared to other networks. In particular, the probability that an outbreak seeded in a single randomly selected individual will spread to at least 5% of the population is biased in this model.

There is a consistency in the values of the epidemic threshold (Figure 1a) and the *R*(*x*; *α*) values for the ensemble of networks generated using the ERGMs (Figure 6). The model “edges” and model “node attributes” networks have higher threshold values than those of model “GWESP = 0.25” and model “GWESP = 0.5”. The *R*(*x*; *α*) plots also point to the same result. It can be seen that the propagation of infection on most model networks is significantly different from that on the original network.

The *R*(*x*; *α*) curves in Figure 6 are also consistent with the epicurves in Figures 2 and 3a. This means that the networks obtained from models “GWESP = 0.25” and “GWESP = 0.5” have a much higher probability of producing an attack rate of at least 5% than school 86 or the others. Figure 4b shows that the overall attack rates for Faux Magnolia are different from those obtained for school 86. The attack rates are calculated for all values of probability of transmission, 0 ≤ *x* ≤ 1. This plot shows that for lower values of *x*, both the networks have similar attack rates, but as *x* increases, they differ. This again agrees with the *R*(*x*; *α*) curves in Figure 4a. The estimated *R*(*x*; *α*) curves for the two networks calculated using the transformed *x* values obtained from the re-calibration method are plotted in Figure 7.

From Figure 8, it can be concluded that the difference in outbreak size on the three networks, Faux Magnolia, school 86 and net1, is not a simple function of time. Figures 8a and 8c show that the epicurves for net1 are very different from the other two. Therefore, for a fixed value of *x*, Faux Magnolia performs much better than net1 in predicting an outbreak on the original school network. Faux Magnolia and school 86 networks produces similar outbreaks in many respects, but there are systematic differences in the duration of the epidemic and the height of the peak values, as shown in Figures 8b and 8d. These differences remain even when the attack rates are matched using the re-calibrated values of *x* as displayed in Figure 8f. Figure 8e leads to a surprising result: for a re-calibrated value of *x*, the less constrained random graph (net1) performs as well as, and sometimes better than, the Faux Magnolia network in estimating the outbreak.

The effects of intervention measures like vaccination are explored by removing nodes at random from the networks. The epicurves before and after the removal of 10 and 100 nodes from the three networks school 86, Faux Magnolia and net1 are shown in Figures 9 and 10 respectively. As expected, the overall attack rate decreases as more nodes are removed for both values of transmission probability, *x* = 0.85 and 0.92. However, Figures 9b, 9d, 10b, 10d and Table 2 show that the magnitude of this effect is different for school 86 and Faux Magnolia. Figure 11, showing the *R*(*x*; *α*) curves for the networks with 10 nodes removed, is consistent with this result.

To verify whether the same re-calibration is valid for the networks with the nodes removed, the transformed value of *x* for Faux Magnolia network is plotted as a function of the *x* for school 86 in Figure 12 for *α* = 0.08. The solid curves show how the effective *x* for Faux Magnolia varies with *x* for school 86. The red curves represent the original networks and the green ones, when 10 nodes are removed. This figure suggests that when the nodes are removed, a different re-calibrated value of *x* would be required to obtain similar attack rates. This is not unexpected. There is no reason to presume *a priori* that the same re-calibrated *x* value would work when the network is changed. Although, it is observed that the same re-calibrated value gives similar results, it is purely coincidental. From Figure 12, it can be concluded for 0.93 ≤ *x* ≤ 1, the original re-calibration is changed very little. For any other values of *x*, there is a high probability that the same re-calibration won’t work so well.

## 4. Conclusions

The transmission of infectious diseases can be investigated as a diffusive process on networks. The topology of the network affects the course of the propagation of the infection through the population. Even on networks with the same degree distribution, number of triangles, clustering coefficients or centrality measures, the course of a disease through a population may vary. Here, the epidemic potential measured by *R*(*x*; *α*) is used to measure dynamically important structural differences between networks. This measure depends on both the global structural aspects of the contact network and the dynamics on the network.

Exponential random graph models are used to generate a number of different networks that match local statistics of one of the friendship networks from the first wave of the Add Health study. The Faux Magnolia network is one such network well known in the literature. Network measures like the epidemic threshold for these two networks are similar, suggesting that Faux Magnolia is a better model for the high school friendship data than others. However, it is observed that there are significant systematic differences in the spread of diseases on the two networks. This implies that the model generating Faux Magnolia does not constrain a set of statistics that is sufficient to reproduce epidemic dynamics.

The epidemic potential *R*(*x*; *α*) for all these networks shows that the school 86 network is more resistant to large outbreaks than any of the others. Treating the transmission probability, *x* as a free parameter, these models can be calibrated so that they all have the same epidemic potential. But the resulting epidemic curves exhibit systematic biases in the peak height and the outbreak duration. Indeed, it turns out that a calibrated, but less constrained, system performs better than Faux Magnolia, suggesting that network re-wiring involved in matching local statistics has introduced spurious global structure.

Moreover, as suspected, different networks do exhibit different responses to interventions. The re-calibration suggested by recognizing the network and the transmission probability are not separately identifiable parameters can be applied to almost any two networks to obtain similar attack rates. There is no *a priori* reason to expect the same re-calibration to be valid after an intervention changes the network structure, even though, as in the networks studied here, the re-calibration may be similar, coincidentally.

It can be concluded that attack rate depends on a mixture of network statistics that goes beyond degree distribution and clustering and is sensitive to some global topological features. The question of what that mixture is requires further investigation.

## Acknowledgement

We thank our external collaborators and members of the Network Dynamics and Simulation Science Laboratory (NDSSL) for their suggestions and comments. This work has been partially supported by Defense Threat Reduction Agency Comprehensive National Incident Management System Contract HDTRA1-11-D-0016-0001 and by the National Institute of General Medical Sciences of the National Institutes of Health under a Models of Infectious Disease Agent Study (MIDAS) Grant 5U01GM070694-13. The content is solely the responsibility of the authors and does not necessarily represent the official views of the National Institutes of Health or the Department of Defense. This research uses data from Add Health, a program project directed by Kathleen Mullan Harris and designed by J. Richard Udry, Peter S. Bearman, and Kathleen Mullan Harris at the University of North Carolina at Chapel Hill, and funded by grant P01-HD31921 from the Eunice Kennedy Shriver National Institute of Child Health and Human Development, with cooperative funding from 23 other federal agencies and foundations. Special acknowledgment is due Ronald R.Rindfuss and Barbara Entwisle for assistance in the original design. Information on how to obtain the Add Health data files is available on the Add Health website (http://www.cpc.unc.edu/addhealth). No direct support was received from grant P01-HD31921 for this analysis.

